# Inactivation effect and damage of multi-irradiance by UVCLED on *Acinetobacter baumannii*

**DOI:** 10.1101/2021.11.19.469185

**Authors:** Mengmeng Li, Baiqin Zhao, Hanlei, Wangzhen Paste

## Abstract

It is acknowledged that the inactivation of ultraviolet has been widely used in various fields. Much literature has been reported that ultraviolet C caused DNA damage to achieve inactivation of microorganisms. There is a lack of unified dose calibration and related parameters in this field. In this study, we used a device consisted of the LED of 272 nm to conduct sterilization experiments against A. baumanii. We confirmed the effectiveness of ultraviolet C sterilization for both sensitive and drug resistance strains and explored the relationship between bactericidal rate and ultraviolet doses under various irradiance. Dose requirements of various irradiance were clarified. High irradiance improved sterilization efficiency greatly. The overall damage to the total genome was observed though gel electrophoresis. Ultrastructure of damaged bacteria were investigated by transmission electron microscope in detail. The study revealed that damage to DNA and to the cytoplasm matrix and ribosomes. The study has yielded the possible effects of ultraviolet light on cells by amplifying the energy. The radiation significantly promoted the production of cell wall and cellular membrane.

**Significance Statement:** The statistical results of bactericidal efficiency are influenced by the quantity of bacteria on the medium. Irradiance on the target surface affects the sterilization efficiency directly. And doses required in low irradiance are much more than high. A high irradiance reduced dosage efficiently to achieve the sterilization which improves the sterilization efficiency. There is a difference between low and high UVC dosage damage to the structure of bacteria. Less energy can make DNA coagulation solidified or be dispersed to the edge. Meanwhile the cytoplasm matrix is ruined. When the energy was enough, there is a boost of cell wall and cellular membrane production. The invisible light causes comprehensive damage to bacteria.

## Introduction

Acinetobacter baumannii has a strong ability to obtain kinds of drug resistance and cloning transmission and now become a worldwide epidemic[1]. It has become one of the most important pathogenic bacteria in nosocomial infections in China. A. baumannii is a conditional pathogen, which causes a variety of diseases such as nosocomial acquired pneumonia and mechanical ventilation associated pneumonia, postoperative and post-traumatic intracranial infection, skin soft tissue infection, etc. However, the main response strategy in the clinic remains hygiene and antibiotic treatment, and it is urgent to find a new effective method for Bactericidal therapy[1, 2]. For the consideration of clinical surgical infection, Acinetobacter baumannii was picked for this study.

People have realized that sunlight has sterilization effect for years which mainly depended on ultraviolet spectrum, especially in the wavelength of 200-280 nm [3, 4]. Ultraviolet light is invisible electromagnetic radiation. According to the World Health Organization, UV is divided into four regions. UVA in the range of 320-400 nm can penetrate into the dermal layer of human skin; UVB at 280-320 nm can reach on the basal layer. While UVC (200-280 nm) can only reach the upper layers and has implications for wound care [5, 6]. And the wavelength of 0-200 nm belongs to vacuum irradiation. UVA is the main factor that induces melanin resulting in skin cancer, and UVB causes erythema damage[7]. The Nucleic acid absorption spectrum indicates that the main absorption of nucleic acids is by UVC in the range of 250-270 nm which does most severe damage to organisms[8-11]. The region strongly absorbed by nucleic acids is most lethal spectrum. And DNA and RNA displayed the highest absorption coefficients of all cellular components. Their main products are cyclobutene-pyrimidine dimers (CPDs) and 6-4 photoproducts (6-4PPs, which are pyrimidine adducts), especially CPDs in DNA. It makes nucleic acid difficult to self-replicate[12-14]. These studies infer that ultraviolet firstly induce DNA and RNA abnormalities and further impair their genetic function so that cells cannot proliferate, and the physiological response is blocked, leading to cells death. However, whether the light cause specific damage to other structures is unclear. Many researches concentrated on the damage to DNA but few phenomena visually showed the effect on cellular structure [7, 15, 16]. In this research, we verified that the genome was not broken into fragments and provided pictures containing intracellular structural destruction information.

Based on the effect of disinfection, ultraviolet rays have been applied in many fields, such as water disinfection system [17], food antiseptic[18]. Researchers have long sought of the application of UV inactivation for infected wound [19, 20]. In 1965, Freytes et al. irradiated patients with oral ulcer though UVC every few weeks and obtained good treatment results[21]. It is the first report on the clinical practice of ultraviolet rays. Over the same period, scientists conducted a large number of experiments though low-pressure mercury lamps to confirm the effectiveness of UVC inactivation on multiple organisms, which consisted of a variety of bacteria, viruses, spores and cysts[22]. Limited by sources of light, the clinical application of ultraviolet C is sluggish. In hospitals, ultraviolet were applied for sterilization in environment such as in operating theatres to obtain the cleanest indoor air[23-25], as well as disinfection of medical devices like catheters[26, 27]. In addition, experiments have shown that ultraviolet light can be used to kill drug-resistant bacteria effectively. Buonanno et al. choose Kr-Br quasi-molecular lasers to emit ultraviolet light at 207nm. They confirmed the wavelength was able to kill MRSA and they further conducted experiments of wound infections on mice[28, 29]. The effectiveness has also been reported by the study which used a Kr-Cl quasi-molecular laser to emit 222nm wavelengths to treat MRSA-infected mice wounds[30].

The traditional low-pressure mercury lamps have wide spectrum which ionizes oxygen to ozone and benefits sterilization but a threaten to health; the quasi-molecular lasers are large in volume with poor regulation and they need to add a filter to guarantee a single wavelength[29]. Both contain toxic elements. In contrast, UVLED emits a single spectral wavelength with no side effects; the size is small which is convenient to assemble and carry. They are inexpensive and durable[31]. In recent years, many scientists have made great progress in semiconductors devices[32]. And the luminous efficacy of UVCLED has been improved [33]. While for medical application, much more researches are needed. Among related researches, most focused on the effectiveness of bactericidal but often ignore the effects of irradiance[34, 35]. Although some researchers have suggested that the irradiance values affected microbial inactivation, few results made a comparison about the difference[36]. And Gora et al. attempted to change the distance to obtain results of different irradiance, restricted by the power of the light, the difference was hard to distinguish[37].

In this study, we designed a light source device with high quality UVCLED. We adjusted the distances between LED and targets to change the irradiance on the targets and carried out experiments against *A. baumannii*. And we irradiated mediums containing different quantities of bacteria. The results provided UV dosages of sterilization rates at 99% 99.9% 99.99% and characterized the sterilization efficiency of different quantities of bacteria. We also confirmed the germicidal efficacy of different irradiances varied greatly. The damaged genome was performed by the electrophoresis and the internal structure of destructed bacteria were observed by transmission electron microscope.

## Materials and Methods

### UVCLED device

The wavelength of UV-LED (BRT-B35CD7A1CSD, High Power Lighting, China) in the study was 272 nm and the divergence angle was 100 °. The LED had no significant thermal effect. The irradiance of the lamp was measured on the face of the targets[38]. The distribution of irradiance of the entire area at same distance was uneven (Fig.1 *B*). We measured at characteristic points and then took the average value (p<0.0001). The irradiance measured by UVCLED Probe (Linksun Technology, China) at the distances of 8.2 4.6 and 1.3 cm was 0.06 0.3 and 1.0 mW/cm^2^ respectively.

**Fig 1.**
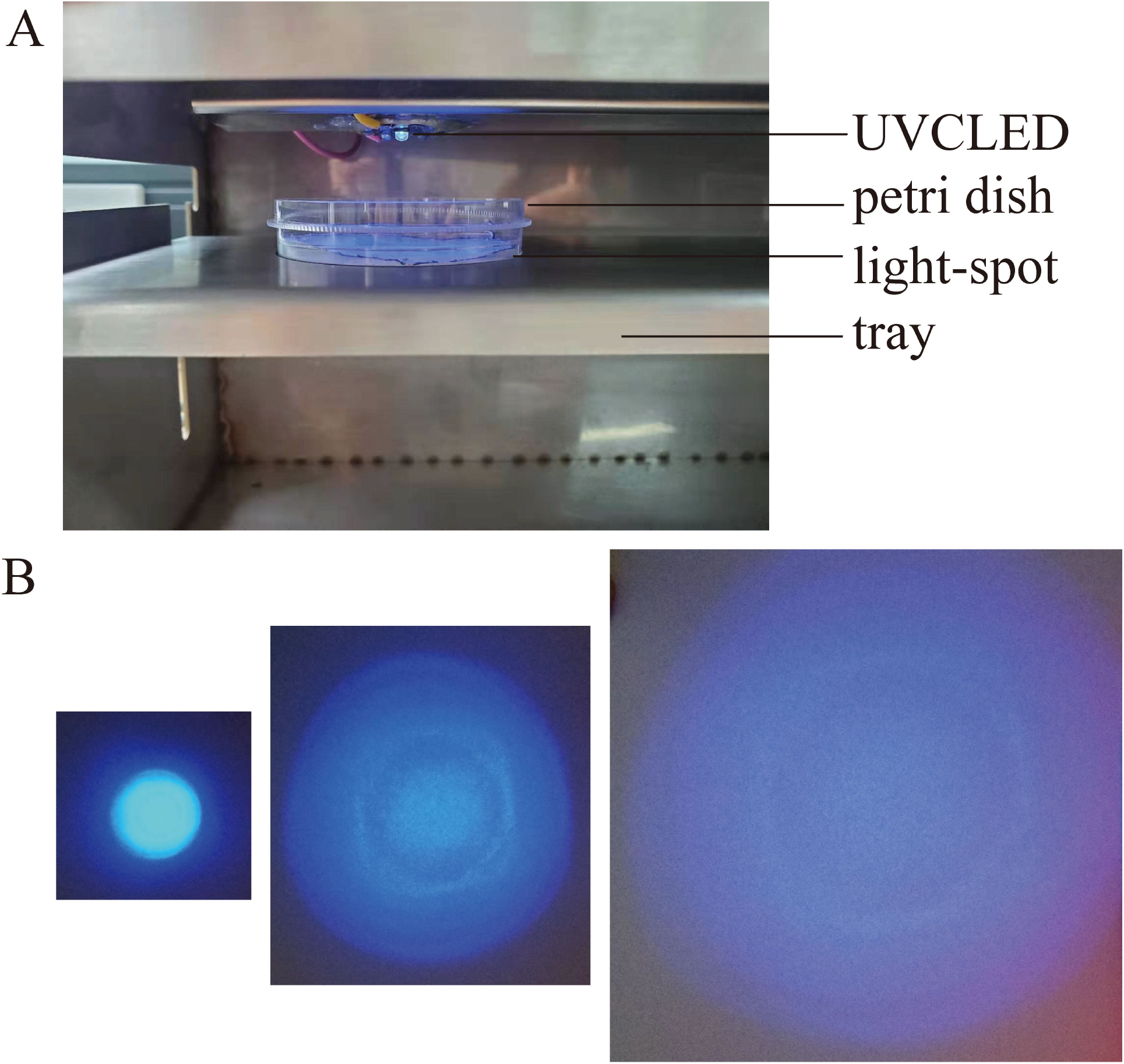
Work scene with the light source device; (A) Example of irradiation scene; (B) Size of light spot of UVCLED at different distances.

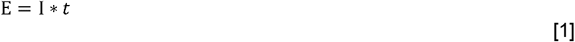

The ultraviolet flux was calculated as the quotation:

Where *E* is UV dosage (mJ/cm^2^), *I* is the irradiance detected on the surface of targets (mW/cm^2^), and *t* is the irradiation time(sec).

There was a time error that we skipped the time for the device from start-up to output, which means, the time for bacteria exposure was a bit less than irradiation time.

### Bacterial Strains, culture Conditions, and Irradiation

For the consideration of clinical surgical infection, Acinetobacter baumannii was used for the experiment. And 5116 was sensitive strain and 32 was multi-drug resistant strain but sensitive to compound sulfamethoxazole specifically. Both were from Xi’an Jiaotong University. They were preserved at minus eighty degrees. To obtain enrichment solution, we picked up single colony by loop and inoculated into 5 mL Luria-Bertani broth and set the glass bottle with suspension in shaking incubator (37°C; 200 rpm). After 14-16 hours, the concentration of bacterial solution was evaluated by the optical density at 600nm measured by ultraviolet spectrophotometer. We collected them by centrifugation (2 min; 10,000 rpm), resuspended in broth and adjusted to an optical density at 600 nm of 0.2.

To get the certain quantities bacteria suitable for irradiation [28]. We took 100μL from the bacterial suspension and carried out ten-fold serial dilutions. Absorbed 100μL in multiple concentrations to spread on agar mediums until they dried out. Put all mediums in a thermostatic incubator at 37°C and observed the growth after 18hours. The number of bacteria colonies on agar medium ranging from 30-300 was effective to count. Those covered properly were prepared for irradiation experiments. Irradiated immediately after coating. Put the petri dishes on the tray and aligned the center of the plate to the LED directly. The digital timer (DH48S-1Z, ECNKO, China) were used to control the time. After exposure, put mediums into the incubator (37 °C) for 12-18 hours. Counted colonies and recorded the images by Umax2000scanner.

### Agarose gel electrophoresis

To explore the UV effect on DNA. Genome of sensitive strain of *A. baumannii* was extracted for electrophoretic experiment[39]. Bacteria untreated were used as control. And the bacteria heated at 100°C for 10 min was the positive control. The bacteria exposed to one min (18 mJ/cm^2^) and 10 min (180 mJ/cm^2^) at 4.6 cm to examine the state of genome. The genome was extracted by Bacteria DNA kit.

The gel making process was carried out in biosafety cabinet. Weighed 0.3 g agarose powder and dissolved in 30ml 1xTAE solution. Heated 3minutes in microwave. After cooling to 60 °C, added 4 μL EB and mixed well, poured into the plastic box, inserted the ruler comb, gently distributed evenly in the plastic box without bubbles. Cooled to room temperature and then put the gel board in a solution filled with 1xTAE, which was higher than the gel. Added marker (5 μL 1000 bp) and the extracted genome samples mixed with loading buffer to the aperture into the hole in sequence. The volume of samples added was calculated according to the concentration to ensure the total amount of the genome was consistent. Covered the electrophoresis tank, connected the circuit, turned on the electrophoresis instrument, adjusted the voltage to 100 V. After 40 min electrophoresis, took out the gel and put it under the gel imager to observe and record.

### Transmission electron microscope

To investigate the effect of UV on bacteria, we observed the inner structure by transmission electron microscope[39]. The bacteria were irradiated with huge doses which were much than the lethal dose requirements. Prepared four mediums spread with 100 μL containing 10^5^ bacteria and cultivated for about 12 hours. One was control, the other samples were taken after treatment by various dosages of ultraviolet C. The experiment procedures were as follows : sample A was exposed to UVC at 8.2cm for 2 hours (432 mJ/cm^2^); sample B was exposed at 1.3 cm for 2 hours (7200 mJ/cm^2^); sample C was firstly diluted in 1500 μL and the liquid was directly exposed at 1.3 cm for 2 hours (7200 mJ/cm^2^). Then we recollected the bacteria separately.

The bacteria were transferred to tubes (1.5 mL) containing 1mL PBS. Centrifuged 4000rpm, 5 min at 4°C. Took out supernatant fluid and added 1 mL 2.5% Glutaraldehyde fixation solution (SCRC, China) gently in 60 s. Don’t blow up the colony. Stored at room temperature for 4 hours.

The following procedures were taken by electron microscope laboratory.

Took off the fixed liquid. Washed the samples by PBS two times. Osmic acid solution (1%) (Johnson Matthey, UK) was used to fixed samples 2 hours at 4 °C. Dehydrated the bacterial samples with ethanol gradients (30–100 %), and then embedded them in the Epon 812 epoxy resin (Structure Probe, USA) at 60 °C for 24 h. The sample was sliced into 70-90 nm slices in ultra-thin slicing machine (LKB, Sweden). The slices were stained with lead citrate solution and 50% ethanol saturated solution of uranyl acetate for 15 minutes, respectively. The thin-section samples were recorded on transmission electron microscope (Hitachi H-7650, Japan).

## Results

### Bacterial distribution density in irradiation experiments

Because the poor penetration of ultraviolet C, the thickness of bacteria layer must be considered. The distribution density of bacteria on the agar medium is the ratio of the number of colonies to the area of the medium. The unit is cells/cm^2^ and the radius is r (petri dish, r = 4.3 cm). Ideally, the solution of bacteria was spread evenly on agar mediums. The size of bacterium is too tiny to observe directly, so we observed the distribution of colonies after 16 hours cultivated on agar mediums to find a proper number of bacteria for exposure experiments. According to the growth (Fig.2 *A*), we determined the initial distribution density at 10^6^ 10^5^ cells/πr^2^ were appropriated for general exposure experiments.

**Figure 2.**
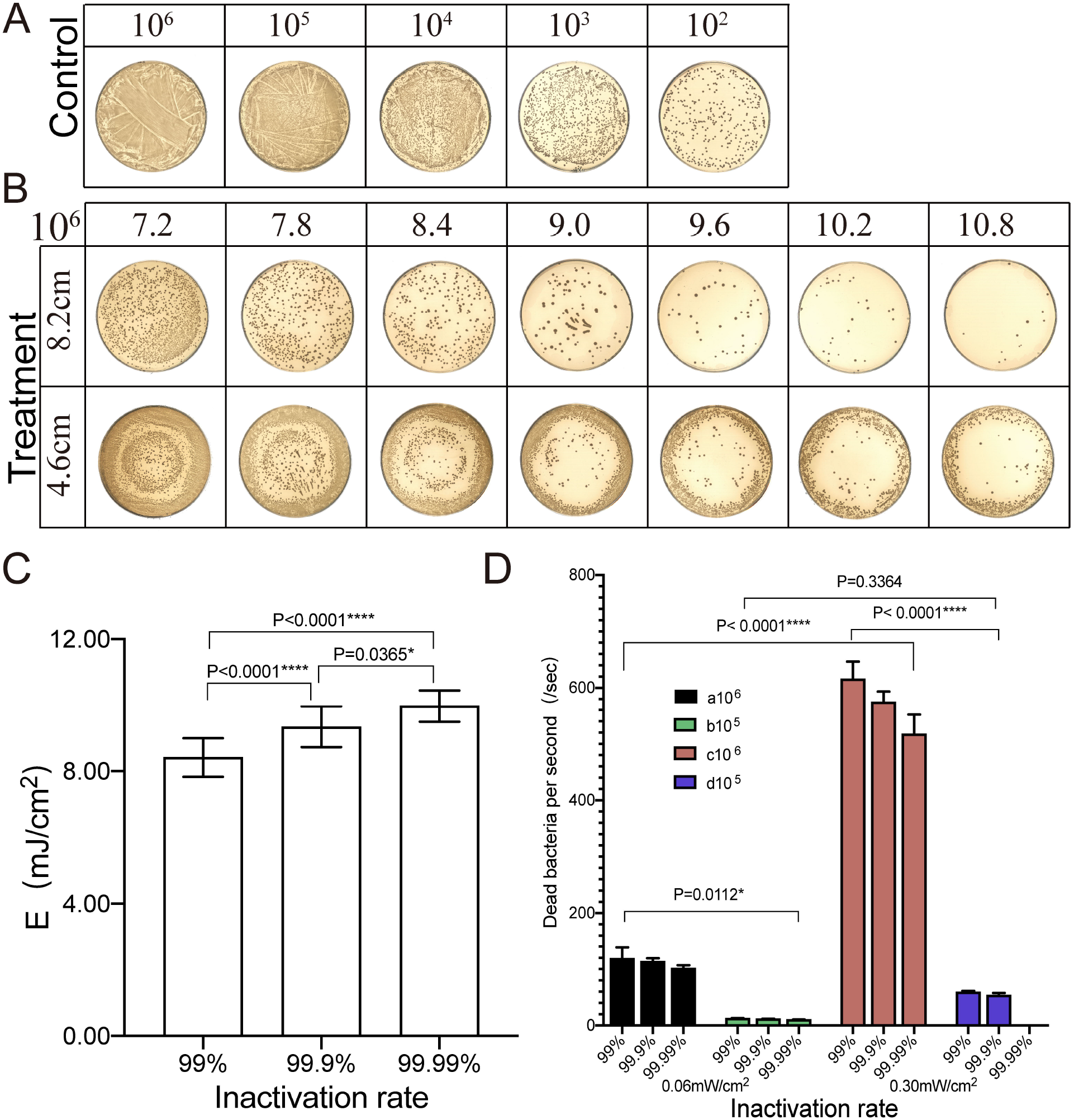
Irradiation results of *A. baumannii* 5116 to achieve different sterilization rates at the distance of 4.6 and 8.2 cm. (A) The distribution of bacteria cultured on agar mediums for 16 hours. The initial bacteria were different. The distribution conditions were as following: dense multi-layer; dense growth; almost monolayer; concentrated; some adhesion; colonies sparsely distributed. (B) The survival of bacteria on agar mediums after exposed at different distances. The initial number was 10^6^. And the doses were consistent. At 8.2 cm, the irradiation time is 120 130 140 150 160 170 180 s; at 4.6 cm, the irradiation time is 24 26 28 30 32 34 36 s. (C) Dose requirements for different inactivation rates. The initial number were 10^5^ and 10^6^ and the irradiance were 0.06 and 0.30 mW/cm^2^. (D) The number of dead bacteria every second. The data was same to (C).

### UV dosage, irradiance, bactericidal rate and efficiency

The sterilization rate is the ratio of the number of colonies that died after exposure to the number of initial colonies[40]. The calculation formula is

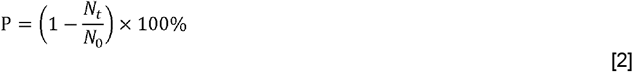

Where N_0_ represents the number of initial bacteria (cells); N_t_ represents the number of colonies grown after irradiation (CFU); t represents the time of exposure(sec) and *P* represents the percentage of bactericidal rate. (1) In the experiment, the ultraviolet spot completely covered agar mediums at 8.2 cm. The counting area was the entire medium. At 4.6cm, the diameter of the radiation area was 6.0 cm, same as counting area. The original quantities of bacteria of both groups were 10^6^. The irradiance at 8.2 and 4.6 cm were 0.06 and 0.30 mW/cm^2^ separately. We kept doses consistent by controlling the irradiation time. All groups showed a gradually decreasing trend of the number of survival bacteria with increasing irradiation time (Fig.2 *B*). It could be deduced that the inactivation effect of ultraviolet C was positively correlated with irradiated doses; that was, the larger the dosages, the more remarkable the effect. And when the energy reached 3.6mJ/cm^2^, the reduction of colonies began to be identified. Furthermore, when the dose reaches 7.8 mJ/cm^2^, single colony counting can be possible; while the dose at 4.6 cm should be 8.4 mJ/cm^2^. (2) To explain the correlation among UV dosage, irradiance, bactericidal rate and efficiency, we set experiments irradiated *A. baumannii* at 0.30 and 0.06 mW/cm^2^. And the order of magnitude of the quantities of irradiated bacteria were 10^6^ and 10^5^. We counted the number of survival though the phenomena of sterilization (Fig.2 *B*). Then we collected all results of four groups for statistical analysis. It suggested that the inactivation effect was positively correlated with the UV dosages (N=7-32, P<0.0001), and for distribution density from 10^6^cells/πr^2^ to 10^5^ cells/πr^2^ and the irradiance from 0.30 to 0.06 mW/cm^2^, bactericidal rates were depended on UV dosages (Fig.2 *C*). And the bactericidal rates at 99% 99.9% 99.99% corresponded to three dose intervals: [8.10, 8.40], [9.00, 9.50], [9.3, 10.80](95% CI of median) respectively. To ensure the effectiveness of inactivation, we took the upper limits of the interval as standard values and the numbers were 8.40 9.50 10.80 mJ/cm^2^ (Upper confidence limit)

Furthermore, to illustrate the differences between four groups, we qualitatively analyzed each experimental condition by comparing any two sets of them. We introduced bactericidal efficiency to characterize the number of dead bacteria per unit time of them at different inactivation rates. The calculation formula is as following,

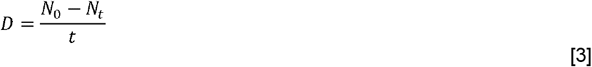

Where *D* represents the quantity of dead bacteria per second (cells/sec); *N*_*0*_ represents the quantity of initial bacteria (cells); *N*_*t*_ represents the quantity of colonies grown after irradiation (CFU); t represents the time of irradiation(sec). There were significant differences in the efficiency under each condition (Fig.2 *D*). Among them, the highest sterilization efficiency appeared in the condition of irradiance at 0.30 mW/cm^2^ and the density of 10^6^ cells/πr^2^; The lowest was in the condition of irradiance at 0.06 mW/cm^2^ and the density of 10^5^ cells/πr^2^. And for each set with same irradiance and densities, the efficiency decreased gradually. When comparing any two sets of data with same sterilization rates and densities, results indicated irradiance at 0.30mW/cm^2^ had higher efficiency than 0.06 mW/cm^2^(Fig.1 *D* ac; bd). For two sets of data with same sterilization rates and irradiance, the efficiency on 10^6^ cells/πr^2^ were higher than 10^5^cells/πr^2^(Fig.2 *D* ab; cd). The results indicated that for irradiance from 0.06 to 0.30 mW/cm^2^, the efficiency was affected by experimental conditions, and the main factor was distribution density. It inferred that the sterilization rates were only determined by UV dosages and ultraviolet C with the irradiance from 0.06 to 0.30 mW/cm^2^ achieved sterilization for bacteria of the density from 10^5^ to 10^6^cells/πr^2^ effectively (Fig.2 *C*). Thereby the inactivation of irradiation on bacteria with larger quantities showed a higher efficiency (Fig.2 *D*). This value was taken to characterize sterilization efficiency of ultraviolet C at certain irradiance. For 0.30mW/cm^2^, the sterilization efficiency were as following: 99%-615.615 cells/s; (SD=31.093, N=2); 99.9%-574.267 cells/s (SD=19.153, N=3); 99.99%-516.980 cells/s (SD=34.998, N=7); For 0.06 mW/cm^2^, 99%-118.529 cells/s (SD=19.973, N=3); 99.9%-113.371 cells/s (SD=6.040, N=11); 99.99%-100.944 cells/s (SD=5.998, N=5). (3) The effect of irradiance on sterilization was further explored by evaluating the phenomena of inactivation at 1.3 cm. The value measured was 1.0 mW/cm^2^. And the spot area was irregular circle, the diameter of the spot area was 2.0 cm while the irradiance beyond that were much lower than this value. When LED was on for four seconds, the sterilization areas appeared and the areas at distribution density of 10^6^ and 10^7^ cells/πr^2^ were 1.9 and 2.0cm in diameter separately. There was obvious boundary between round sterilization area and surrounding area which was covered by bacterial colonies. Sometimes several residuals remained in this area and the sterilization rates were larger than 99.99%. The fields extended slowly with the increase of time. Exposed to equal time, the sterilization areas with low density were slightly bigger than the high, sometimes the same (Fig.3 *A* and *B*). The increase of irradiated areas was nonlinear, indicating that the efficiency was different in time periods. And the two groups have the same trend. To explore the phenomenon, a parameter v was introduced to quantify speed rates of sterilization in different intervals. The calculation formula is as following,

**Figure 3.**
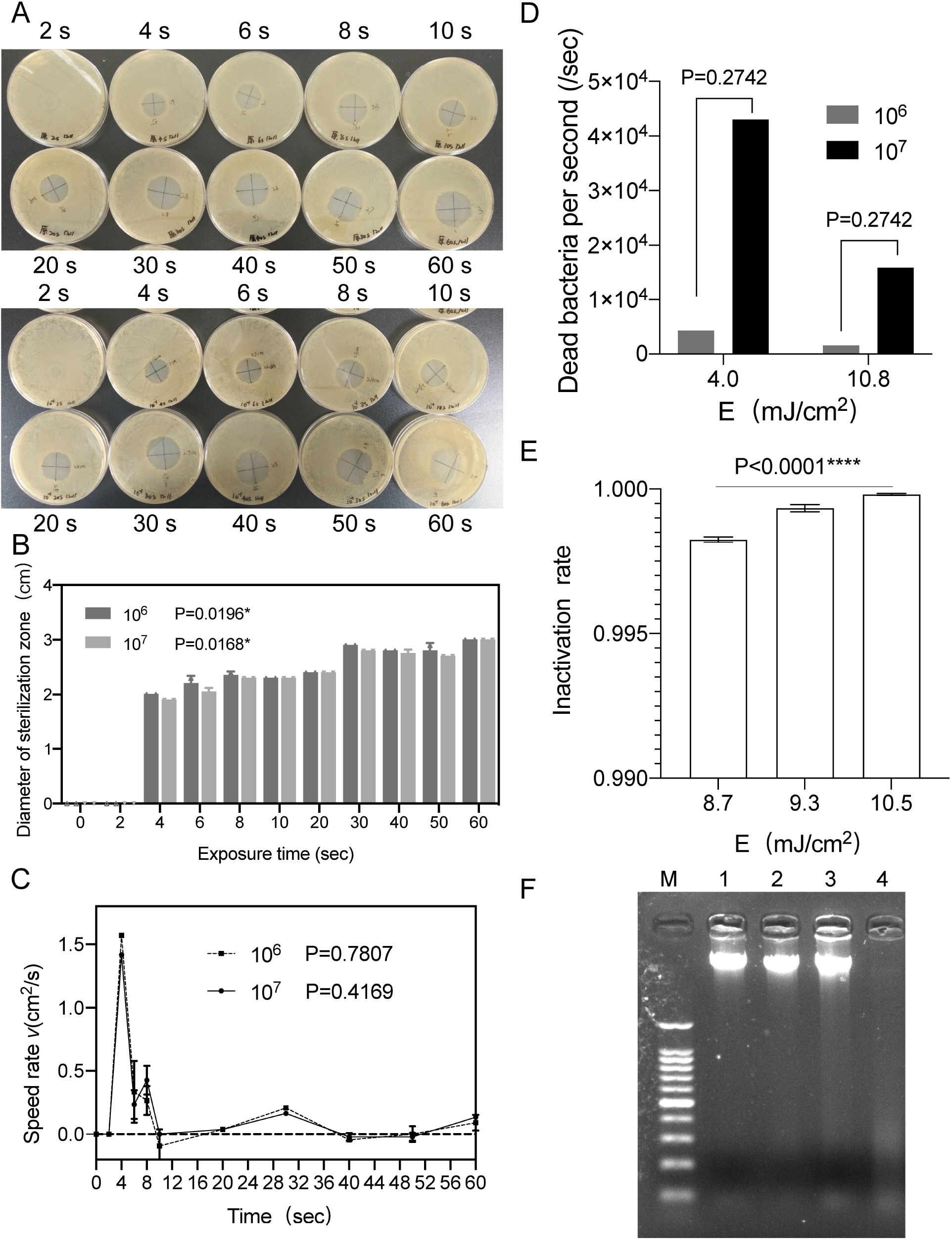
Irradiation results of bacteria at the distance of 1.3 cm; the results of *A. baumannii* 32 and an electrophoresis image of the genome of *A. baumannii* 5116. (A) Pictures of sterilization of bacteria at 1.3cm, the initial number was10^7^ and 10^6^; (B) The changes of diameter of bacteriostatic areas with increasing time for quantities of 10^6^ and 10^7^. (C) The changes of bactericidal speed rate over different time periods. The maximum was at 0-4 seconds, after which the rate decreased sharply and kept steadily. (D) Comparison of two sterilization methods. The dose of 4 mJ/cm^2^ was actual value and 10.80 mJ/cm^2^ was theoretical value. (E) Dose requirements of multi-drug resistance strain at 8.2 cm. (F) Agarose gel electrophoresis of genomic DNA extracted from *A. baumannii* 5116. Lane M: DNA size maker. Lane 1: Without treatment. Lane 2: Irradiated at 4.6 cm for one minute (18 mJ/cm^2^). Lane 3: Irradiated at 4.6 cm for 10 minutes (180 mJ/cm^2^). Lane 4: Heated at 100°C for 10 minutes.

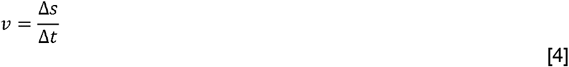

Where v represents speed rates(cm^2^/s); Δ*s* represents areas increased between two irradiation times (cm^2^); and Δ*t* represents time intervals between two irradiation times (sec).

And the peak rate occurred in first interval of four seconds. After that, there was a sharp decline. And with prolonging of exposure, the speed rates of sterilization slowed down and kept stable (Fig.3 *C*). The areas extended after four seconds resulted from the accumulation of dose, which displayed a low speed of inactivation. The results indicated that there were two kinds of inactivation, one reached sterilization rate at 99.99% in 4.0 mJ/cm^2^, another took much more time to accumulate energy to about 10.80 mJ/cm^2^ or much more to achieve sterilization. It suggested that the different irradiance caused various damage to bacteria and high irradiance caused severer destruction rapidly than low. Higher irradiance significantly reduced energy consumption.

### Lethal effect on antibiotic resistance strain

Previous experiments had confirmed the inactivation of sensitive strain. To obtain the result of resistant strain, we conducted experiments with *A. baumannii* 32, a multi-drug resistant strain. The mediums contained 10^6^ cells were irradiated 145 155 175 seconds at 8.2 cm separately, at which the doses were 8.7 9.3 10.5 mJ/cm^2^. The corresponding bactericidal rates were 99.82% 99.93% and 99.98% (Fig.3 E). Compared to the sensitive strain, the results showed their coincidence was well in uncertainty range. It inferred that ultraviolet C was effectively against drug-resistant strains as well.

### Agarose gel electrophoresis

In this experiment, UV effect on DNA was investigated by electrophoresis experiments on extracted genome from the sensitive strain. We compared the migration of samples in the gel to estimate the damage induced by ultraviolet C (Fig.3 F). The ladder of marker showed the fragments lengths it contained. And total genes extracted had no segments separated. It was obvious that the control was a state of genome-reunion and a little away from the loading hole without any isolated fragments. There was no difference between the samples irradiated by 18 mJ/cm^2^ and control. While the sample exposed to 180 mJ/cm^2^ indicated the segments were not completely reunited with some scattered. And as the positive control, the gene molecules after high temperature had disappeared totally.

### Transmission electron microscope observation

To further explore the effect of ultraviolet C, we observed internal structure of *A*.*baumannii* treated with various UV doses though transmission electron microscope. A normal bacterium was coccobacillus morphology with well-formed complete structures. The contours of cell wall and cellular membrane was clear. The dark gray colloidal substance was cytoplasm. The irregular black areas in cells were nucleoids containing DNA and the dark particles distributed in the cytoplasm were ribosomes. No capsule, no flagella. There were white gaps in the background. Principally because the bacteria had thick cell wall and dense structure which made the fixed solution difficult to permeate through. So that the film was not firm and easy to be torn and formed pores. The picture showed cells at different stages, such as individual cells and cells those were dividing (Fig.4 *A*). For the bacteria exposed by 432 mJ/cm^2^, the number of white gaps decreased. And the light transmittance of the film increased. Although the bacteria maintained their shapes, the cytoplasm became transparent while the edge darkened. The viscosity of cytoplasmic matrix decreased. Furthermore, the DNA aggression became vague and dispersed to the edge of the bacterium. The same phenomenon occurred in bacterium that was dividing (Fig.4 *B*). For the bacteria exposed by 7200 mJ/cm^2^, most remained the shape, and there were two different forms of internal damage. One was that the substance in nucleoid gathered into clusters, and the cell wall and cellular membrane thickened. Another was that DNA dispersed to the cell edge and the cytoplasm became more transparent. Some clumps of DNA escaped from the cells (Fig.4 *C*). It was clear that the most damaged cells were those irradiated in liquid. The fixative fluid completely penetrated into the cells and there was high light transmission with no gaps in background. The structures of most cells were incomplete and cell debris scattered. Some kept intact but cell volumes swelled about four-fold. The pictures exhibited that the cellular components were completely destructed and that the gel state of cytoplasm disappeared where aggregates were mass scattered or concentrated distribution aggregated inside the bacteria. The cell wall thickened significantly. And there was intracellular fluid exuded from cells. Some bacteria had gaps in cell walls. And for some bacteria, the cytoplasm was partly isolated though the cell wall (Fig.4 *D*).

**Figure 4.**
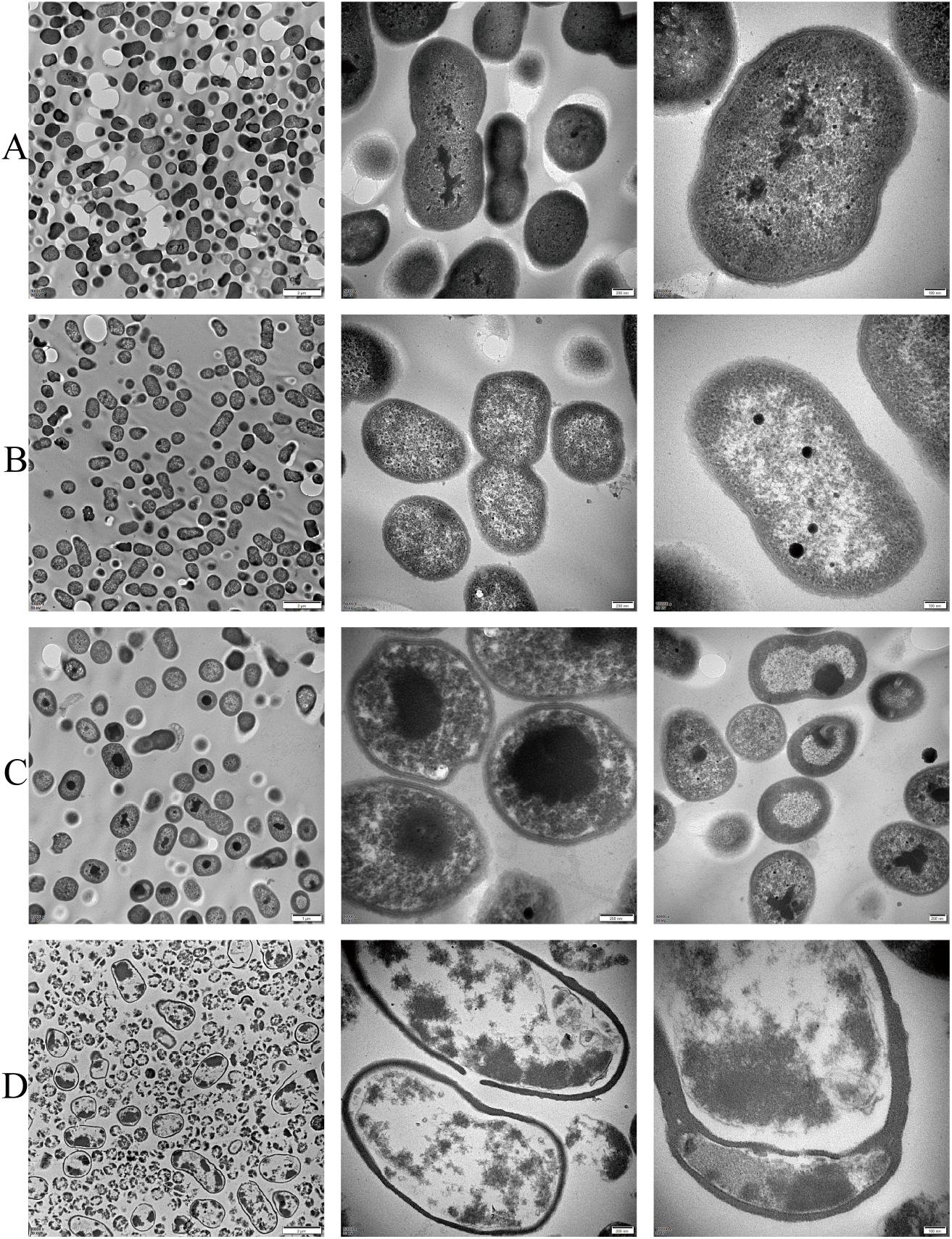
Transmission electron microscopy images of *A. baumannii* 5116. The magnification increases from left to right at 80 kV. And the magnification was different. (A) Normal morphology without treatment. large number bacteria; dividing bacterium; inner formation of bacterium. (B) Irradiated at 8.2 cm for 2 hours (438 mJ/cm^2^); damaged bacteria; dividing bacterium; inner formation of bacterium. (C) Irradiated at 1.3 cm for 2 hours (7200 mJ/cm^2^); damaged to varying degrees; DNA coagulation; DNA shrunk or dispersed. (D) Irradiated in liquid at 1.3 cm for 2 hours (7200 mJ/cm^2^); damaged bacteria; bacteria with intact or ruptured cell walls; bacterium with layered cell wall and membrane.

## Discussion

Bacterial infection has always been an important problem in clinical practice, and abuse of antibiotics for years has been contributed to the emergency of drug-resistant bacteria. There is an urge for a new sterilization method. Ultraviolet C is often used for environmental sterilization but less application in therapy. To provide more reliable references, we designed a light source apparatus to clarify the effects of irradiance on sterilization and observed the morphological changes to explore the damage principle.

### Bacterial distribution density in irradiation experiments

As a reference for the invisible, we took pictures of the visible light of the LED to provide a direct-viewing distribution of energy (Fig.1 *B*). In general, the beam energy emitted by the LED was uneven and roughly decreased gradually from the center to the edge in a plane. The picture showed it contained a transient increase in the process of reduction. And the distribution of survival bacteria after irradiation presented consistent phenomena (Fig.2 *B*). The bacteria cultured 16hours after irradiated were evenly distributed on the surface of agar mediums. At 8.2 cm, the exposure area fully covered the whole mediums, the quantities of colonies basically decreased from the middle to the edge. While the phenomena at 4.6 cm exhibited that the colonies gradually increased from less and then decreased. It was more remarkable when the doses were below 9.0 mJ/cm^2^. It indicated that when the light spots were small than the target surface, the uneven irradiance exhibited the phenomenon of disinfection. And the results also suggested that the irradiance distribution was decreasing roughly from the middle to the edge with a slight increase.

### UV dosage, irradiance, bactericidal rate and efficiency

In the study, when the irradiance was 0.06 and 0.30 mW/cm^2^, ultraviolet C had the equal bactericidal effect on the same distribution of bacteria and was positively correlated with dosages (Fig.2 *C*). And the effect of irradiance on germicidal efficacy was given by three distance experiments. The results suggested when the irradiance ranged from 0.06 to 0.3 mW/cm^2^, for 10^5^-10^6^ bacteria on the medium, the sterilization rates were depended on the total energy not related to irradiance. It took times to reach the same energy so that the speed rates were different for two irradiances. As the sterilization rate increased, the speed rates decreased (Fig.2 *D*). There were two possible reasons. On the one hand, with the increase of sterilization rate, the number of surviving bacteria gradually dropped, the density descended. But the surface was full covered by ultraviolet C. The light cannot locate the living bacteria automatically but continuously irradiated all bacteria with quantities of dead bacteria, which cause a surplus of energy. On the other hand, the unavoidable condition was that the stacked cells made it hard for ultraviolet light penetrate one cell to another which needed to accumulate much more energy. Both cases prompted energy loss resulting in a reduction of efficiency.

However, the demonstration analysis was inapplicable to the results obtained in the experiments at 1.3 cm. At the irradiance of 1.0 mW/cm^2^, it took 4.0 mJ/cm^2^ to achieve the inactivation rate over 99.99% (Fig.3 *A*). The irradiance of applicable condition for dose requirements was restricted to 0.06-0.03 mW/cm^2^. When it came to 1.0mW/cm^2^, bactericidal effect no longer obeyed the dose requirements. The phenomenon implied that high irradiance dramatically amplified the germicidal efficacy. The sterilization rate of the irradiated area all exceeded 99.99% at 1.3cm and the inactivation speed of multi-irradiance was characterized by the increased sterilization area (Fig.3 *B*). It showed the sterilization efficiency of various irradiance of the spot had great differences. The correlation is non-linear ((Fig.3 *C*, R^2^ < 0.1).

If the radiation intensity on the entire medium was 1.0mW/cm^2^, the time to reach 99.99% must be 4 seconds. Therefore, we can compare the following scenarios. Under the condition that the number of bacteria is 10^6^, we set another condition that the irradiation covers the whole medium with irradiance at 1.0 mW/cm^2^. Assuming the irradiance conforms to dose-dependence just as 0.06-0.3mW/cm^2^, the dose requirement for 99.99% should be 10.80mJ/cm^2^ though 10.80seconds. And the speed rate is supposed to be 5.38 cm^2^/s (equation [4]). In fact, the bactericidal time demanded is 4 seconds (Fig. 3 *A*), and thus the corresponding speed rate should be 14.51cm^2^/s (equation [4]) which is 2.70 times the hypothetical data. And for the inactivation efficiency, the actual value of the quantities of dead bacteria was 2.70 times (equation [3]) the previous (Fig. 3 *E*).

The counterevidence demonstrated that the energy of irradiance at 1.0mW/cm^2^ did significantly improve the sterilization efficiency. For the irradiance it was unnecessary to meet the former dose demand. But for the irradiance much lower than 0.06 mW/cm^2^, the dose demands may be larger. Consequently, to achieve UV sterilization must attain a certain dose but demands were diverse due to the varied ability of irradiance. The dose requirements were united within the range of 0.06-0.3 mW/cm^2^ but were reduced substantially at 1.0 mW/cm^2^. Irradiance effect could be considered in practical application.

### Lethal effect on antibiotic resistance strain

The results of both sensitive and multi-drug resistance strain at 0.06 mW/cm^2^ suggested that two strains had equal tolerance to ultraviolet which proved that the wavelength effectively affected the drug resistant bacteria. More importantly, it inferred that the dose requirements for sensitive strain were appropriated to antibiotic-resistant strains as well (Fig.1 *C* and Fig.2 *E*). The genomes of drug-resistant strains with special genes made them against antibiotics which distinguished them from the sensitive. But the effect of ultraviolet was not specific to some drug-resistant genes. Resistance had no obvious effect on ultraviolet sterilization. It could be predicted that UV sterilization to all strains was consistent with a little difference in growth ability.

### Agarose gel electrophoresis

The gel electrophoresis image partly displayed the injury of DNA from *A. baumannii* after treated by ultraviolet C. Unlike the destruction by high temperature, the doses supplied did not break the genome into small segments directly. And the strips of two treated samples suggested that higher energy caused severer damage. The dose of 180mJ/cm^2^ made the genome unstable and disperse under the traction of the charge (Fig.3 *F*). This damage further influenced the replication and transcription of DNA, leading to cell death. And the concrete damage to DNA structure was still unclear.

### Transmission electron microscope observation

The transmission images of *A. baumannii* provided pictures containing internal ultrastructure in detail which clearly revealed the damage of ultraviolet radiation to bacteria (Fig.4). The different shapes were shown in the figure because the samples were randomly sliced, which were transversely cut into spherical and longitudinally cut into long oval shapes. Four sets of images showed that ultraviolet light acted on cells in all states, including dividing cells. The irradiated colonies on the surface of medium showed that as the dose increased, the dispersion of the materials in cytoplasm strengthened which provided a distinct boundary between transparent cytoplasm and gray black edge. The bacteria in a small droplet (1mm thick) suffered the greatest destruction that all cellular components were abnormal. Although exposed to equal dosage, for each bacterium, those distributed in liquid had more access to the light directly.

The study showed that ultraviolet C had a variety of injuries to bacteria. Amounts of energy damaged the bacteria deeply. The light turned DNA into flocs or pellets. The colloid of the cytoplasmic matrix was diluted gradually. Permeability of cell wall and membrane increased and allowed external fluid penetrated into. And the liquid inside squeezed the cell membrane and the cell wall which expanded the cell and caused ruptures. As the irradiation time increased, the various parts of cell were distinguished clearly. Another obvious phenomenon was the cell wall thickened. When DNA and cytoplasm were damaged where the physiological and biochemical reactions were affected, bacteria had been increasing genes expression to build up cell walls and membranes. And it had been demonstrated that the regulatory factors of membrane and wall formation were not destroyed by ultraviolet light which inferred that the energy stimulated the expression of some genes to thicken cell wall and membrane. The visible defense of bacteria against ultraviolet C was the intensification of cell wall and cellular membrane.

## Conclusion

In the experiment, the UVCLED device was well applied for various irradiation conditions. And we explored the relationships among irradiance, dosage, efficiency and bactericidal rate in detail by the multi-irradiance sterilization experiments. We also verified the efficacy of UVC on drug-resistant strain. And damaged DNA was examined by electrophoresis experiments. The internal structures were observed through the transmission electron telescope. The results showed that the sterilization rates were only positively correlated with the irradiated doses at the irradiance of 0.06-0.30 mW/cm^2^ and when irradiance was greater than 1.0 mW/cm^2^, less dose achieved sterilization rate. And inactivation effect of ultraviolet C showed no selectivity for drug-resistant strain, which is conducive to the promotion of UVCLED application. However, whether high energy bring more serious damage to wound is still under research. UV effect on bacteria first manifested in the abnormality of DNA, as dose increased, the accumulated energy exhibited a destructive effect on other structures. And ultraviolet C made the nucleoid shrink and destroyed the cytoplasm of the physiological biochemical reaction site but didn’t affect the pathway of cell wall and cell membrane production and enhanced the formation of them.

## Abbreviations

UVCLED: ultraviolet C light emitting diode
CFU: colony-forming units
*A.baumannii*: *Acinetobacter baumannii*
PBS: phosphate buffer

## Acknowledgments

The work was supported by the Institute of semiconductors, Chinese Academy of Sciences.

Thanks to Xi’an Jiaotong University.

No uncommon reagents or instruments.

The light device is protected by patent. Patent application number: bdt2021-342

## Notes

### Competing Interest Statement

Baiqin Zhao designed the study, LeiHan provided some experimental techniques, and Zhen Wang helped in Light device and Mengmeng Li conducted the research, collected and analyzed data and wrote this article. No uncommon reagents or instruments. The light device is protected by patent. Patent application number: bdt2021-342

